# Modeling of root nitrate responses suggests preferential foraging arises from the integration of demand, supply and local presence signals

**DOI:** 10.1101/2019.12.19.882548

**Authors:** Meine D. Boer, Joana Santos Teixeira, Kirsten H. ten Tusscher

**Affiliations:** Computational Developmental Biology Group, Utrecht University, The Netherlands

**Keywords:** phenotypic plasticity, preferential root nitrate foraging, modeling, demand and supply signals

## Abstract

A plants’ fitness to a large extent depends on its capacity to adapt to spatio-temporally varying environmental conditions. One such environmental condition to which plants display extensive phenotypic plasticity is soil nitrate levels and patterns. In response to heterogeneous nitrate distribution, plants show a preferential foraging response, enhancing root growth in high nitrate patches and repressing root growth in low nitrate locations beyond a level that can be explained from local nitrate sensing. Although various molecular players involved in this preferential foraging behavior have been identified, how these together shape root system adaptation has remained unresolved. Here we use a simple modeling approach in which we incrementally incorporate the various known molecular pathways to investigate the combination of regulatory mechanisms that underly preferential root nitrate foraging. Our model suggests that instead of a thus far not discovered growth suppressing supply signal, growth reduction on the low nitrate side may simply arise from a reduced root foraging and increased competition for carbon. Additionally, our work suggests that the long distance CK signaling involved in root growth increase in high nitrate patches may represent a supply signal specifically functioning in modulating demand signaling strength. We illustrate how this integration of demand and supply signals prevents excessive preferential foraging under conditions in which demand is not met by sufficient supply and a more generic foraging in search of nitrate should be maintained.

## 1 Introduction

Phenotypic plasticity is of critical importance for sessile plants to adapt to and survive in a variable, heterogeneous environment. One of the environmental factors to which the root system of plants display extensive phenotypic variation is soil nitrate availability. Adaptation to spatio-temporally variable nitrate availability entails changes in nitrate storage and assimilation, adjustment in the spatial patterns, types, numbers and affinity of expressed nitrate transporters as well as extensive adjustment of overall root system architecture (RSA) (Aibara and Miwa, 2014). Harnessing the full range of this plasticity may reduce the demand for artificial fertilizers and improve agriculture on poor soils, yet requires an improved understanding of the processes underlying this plasticity. This improved understanding is also needed to help combat the deleterious effects of excess nitrate deposition on natural ecosystem diversity.

Because of the extensive effects of nitrate on root system architecture, nitrate has been termed an environmental morphogen (Guan, 2017), and substantial research has been devoted to unraveling the mechanisms through which nitrate affects RSA. The picture that has emerged is that plants employ a highly complex molecular network responsible for the sensing of internal and external nitrate status, integration of these signals, and the mounting of a suite of possible growth responses. For plants exposed to homogeneous external nitrate conditions, depending on external and hence internal nitrate levels a continuum of growth responses has been described (Giehl and Wirén, 2014). For very low internal nitrate status, plants engage in a survival response, repressing root growth through the CLE-CLV1 module (Araya et al., 2014). The low nitrate induced repression of the AUX/IAA ACR4/AXR5 further contributes to this survival response (Giehl et al., 2014) (Fig 1A, left). For somewhat less low internal nitrate levels plants instead display a foraging response, promoting root growth via the induction of TAR2 which results in enhanced local auxin biosynthesis (Ma et al., 2014). Additionally, this foraging response likely involves the low nitrate status induced expression of WAK4 and the downstream auxin transporter MDR4/PGP4 (Giehl et al., 2014) known to promote lateral root formation (Lally et al., 2001; Terasaka et al., 2005) (Fig 1A, second from left). Finally, for very high systemic nitrate levels, a systemic repression response occurs, repressing root growth through repression of auxin sensing via the AFB3, NAC4 and OBP4 pathway (Vidal et al., 2010). Root growth may be further repressed through the HNI9 dependent repression of nitrate transport (Girin et al., 2010) (Fig 1A, right).

**Figure 1.**
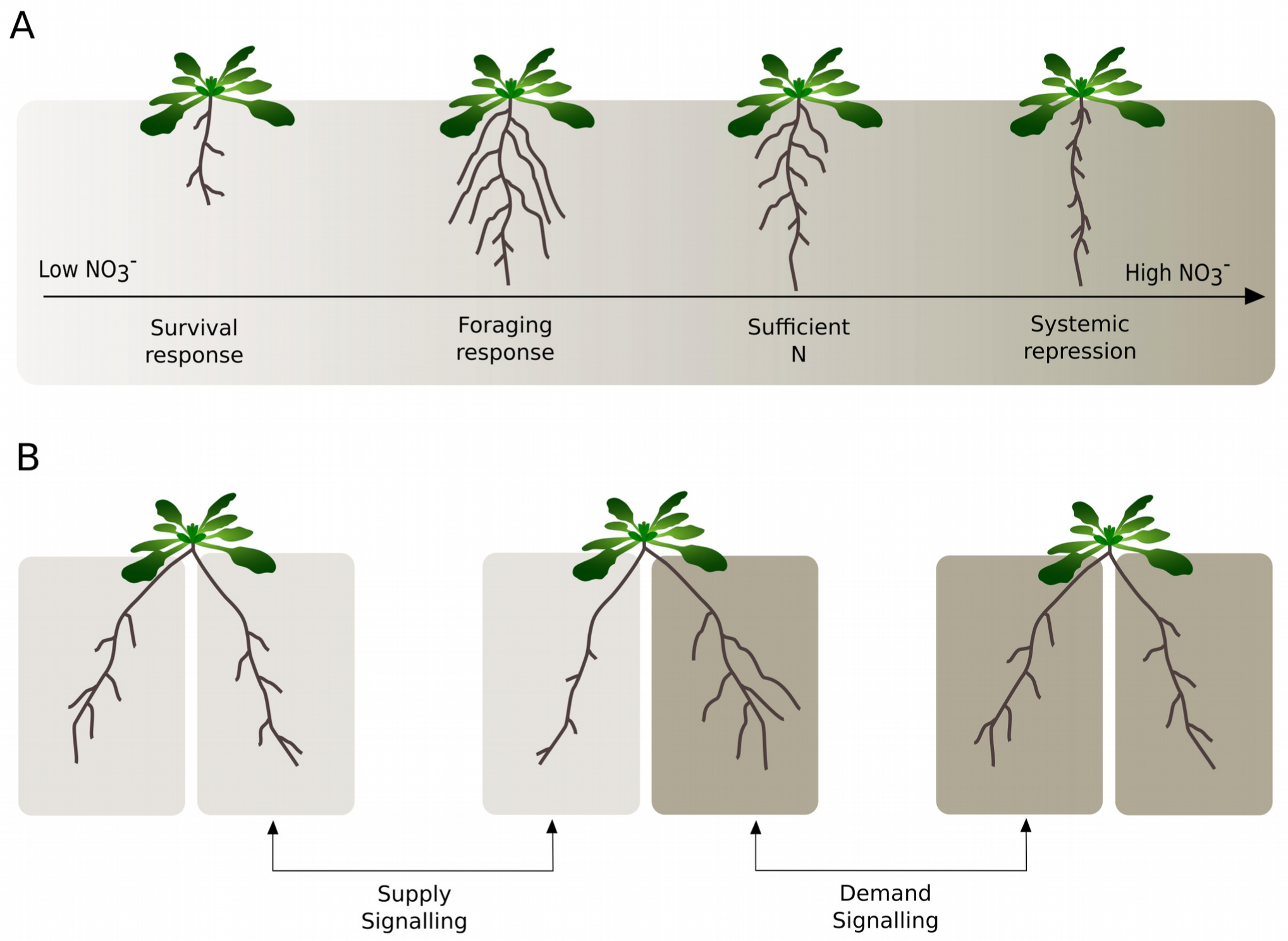
Response of root system architecture to environmental and internal nitrate status. **(A)** Range of growth responses occurring for increasing homogeneous external nitrate levels. For the lowest nitrate levels a growth repressing survival response occurs, somewhat less low nitrate levels induce a growth promoting foraging response, whereas very high nitrate levels induce a growth reducing systemic repression response. **(B)** Response of root growth in a split root architecture to homogeneous or heterogeneous nitrate conditions. Enhanced root growth at the high nitrate side under heterogeneous as compared to homogeneous conditions is indicative of the presence of a growth promoting demand signal. Similarly, reduced growth at the high nitrate side under heterogeneous as compared to homogeneous conditions indicates a growth repressing supply signal.

In addition to these overall RSA growth responses, under heterogeneous external nitrate conditions plant roots display a preferential growth of the root system in nitrate rich patches, a phenomenon referred to as preferential root foraging (Ruffel et al., 2011; Guan et al., 2014; Mounier et al., 2014). To investigate the mechanisms underlying preferential root foraging, split root experiments are used (Ruffel et al., 2011; Guan et al., 2014; Mounier et al., 2014) (Fig 1B). For sufficiently different high and low nitrate levels, a typical outcome of these experiments is that exposure to different nitrate concentrations results in substantial growth asymmetry (Fig 1B, middle). Furthermore, growth at the high nitrate side is enhanced beyond that of a plant experiencing high nitrate at both root halves, whereas growth at the low nitrate side is diminished beyond that of a root system experiencing low nitrate at both sides. These differences have been taken as evidence for the presence of growth promoting systemic demand signals and growth repressing systemic supply signals (Ruffel et al., 2011; Guan et al., 2014; Mounier et al., 2014).

Over the last years, several key players in the preferential foraging of roots for nitrate have been discovered. First, it was found that the dual-affinity nitrate transporter NRT1.1 at low external nitrate levels imports auxin, effectively reducing lateral root auxin levels and thus repressing lateral root growth (Krouk et al., 2010). At sufficiently high nitrate levels, no auxin transport and hence no repression occurs. Instead NRT1.1, via its transport of nitrate, enhances auxin signalling through the AFB3, NAC4 and OBP4 pathway (Vidal et al., 2010, 2013, 2014) which was recently extended with the cell wall modifying enzyme XTH9 (Xu and Cai, 2019), and via its sensing of nitrate enhances auxin signalling via ANR1 (Remans et al., 2006), thereby promoting lateral root growth. Mutation of NRT1.1 severely reduces the preferential root foraging response (Remans et al., 2006; Mounier et al., 2014). A second cornerstone of preferential nitrate foraging was discovered with the elucidation of the C-terminally encoded peptide (CEP) demand signalling pathway. Under low external nitrate conditions, roots locally produce CEP peptides (Tabata et al., 2014), which become translocated to the shoot via the xylem. In the shoot they bind to so-called CEP receptors (CEPR) (Tabata et al., 2014), resulting in the production of downstream signals (CEPD) (Ohkubo et al., 2017). These downstream signals travel back to the root via the phloem, upregulating the nitrate transporter NRT2.1 only in roots perceiving sufficiently high external nitrate (Tabata et al., 2014). NRT2.1 has been shown to be an important component in nitrate dependent root growth stimulation (Little et al., 2005; Remans et al., 2006; Naz et al., 2019). While the mechanism through which NRT2.1 promotes root growth remain to be fully elucidated, in rice it has been shown that nitrate uptake leads to the production of NO, which subsequently results in elevated auxin levels via the upregulation of the auxin transporter PIN1 (Sun et al., 2018). As a consequence auxin supply to the lateral root is increased, enhancing lateral root growth. Thus, upregulation of NRT2.1 appears to also stimulate lateral root growth in the presence of nitrate through auxin, but via a different, less direct mechanism. Finally, an important role for systemic CK signaling in preferential nitrate foraging has been recently uncovered (Ruffel et al., 2011; Poitout et al., 2018). Plant roots were found to produce CK in a nitrate dependent manner, with this CK subsequently being transported to the shoot, where it controls the expression of a large number of genes as well as impacts preferential root foraging.

Intriguingly, mutations in NRT1.1, NRT2.1, CK biosynthesis and CK transport all strongly reduce preferential root foraging or uptake (Cerezo et al., 2001; Ruffel et al., 2011; Mounier et al., 2014), suggesting an integrated, synergistic response network rather than mere additive actions. Still, how exactly these different pathways are integrated and whether their combination is necessary and sufficient to explain the outcomes of split root nitrate foraging experiments remains unclear. First, while split root experiments are generally taken to indicate the presence of growth stimulating demand and growth suppressing supply signals, thus far no supply signal specific for heterogeneous nitrate conditions has been proposed. Additionally, while the role for systemic nitrate levels in modulating root growth dynamics was recently further substantiated (Okamoto et al., 2019), how systemic survival, foraging and repression responses may be involved in generating root growth asymmetry under heterogeneous nitrate conditions has not been studied. Finally, plant organs are in a continuous competition for carbon resources, a process that may contribute to growth asymmetries but which involvement in preferential root foraging so far has not been considered.

In addition to the question which of these mechanisms are involved and how they are integrated to generate preferential foraging, an open question is the spatial tunability of this preferential foraging response. It has been shown that the extent of preferential foraging depends on the concentration differences between high and low nitrate patches, as well as the average nitrate level and hence plant nitrate status (Mounier et al., 2014). Larger concentration differences and lower average nitrate levels have been shown to elicit a stronger foraging response. This can be understood from the fact that in these situations more was to be gained from the high nitrate side as well as that the plant was more in need of nitrate. What is less clear is how the size or number of nitrate rich versus nitrate poor patches impacts the foraging response. As an example, if only a single high nitrate patch has been found by the root system, excessive local proliferation in that patch may go at the cost of root growth elsewhere, prohibiting the discovery of new nitrate rich patches. In situations with only a small part of the root system experiencing high nitrate levels, it may thus be more optimal to keep preferential proliferation somewhat bounded to maintain a minimum level of explorative growth in the rest of the root system.

Here we will use a modeling approach to shed light on the above questions. To this aim we develop a first, simple model for the preferential foraging of roots in nitrate rich patches. We incrementally incorporate known regulatory mechanisms involved in adapting RSA to environmental and internal, systemic nitrate conditions into our model. Following this approach, we identify the likely involvement of systemic nitrate dependent suppression and foraging responses, as well as competition for carbon resources in preferential root foraging behavior. Finally, we propose a novel hypothesis for the role of long distance CK signalling in preferential root foraging, suggesting that it entails a nitrate supply signal that modulates demand signal strength.

## 2 Materials and Methods

### 2.1 Model equations

#### 2.1.1 A basic model for root system growth and internal nitrate dynamics

Our goal is to construct a simple root growth model enabling us to investigate the combined regulatory effects of external and internal nitrate status on root system growth and how these generate preferential foraging in nitrate rich patches. For simplicity, we will describe growth dynamics of (a part of) the root system in terms of changes in its cumulative length *L* (in mm), thus ignoring growth induced changes in root diameter, branching, or differences in growth dynamics between main and lateral roots. As a further simplification, we do not explicitly model shoot growth and the dependence of root growth on shoot generated photosynthesis products. Instead, we assume that shoot leaf area is proportional to root system length, thus ignoring potential changes in root shoot ratio. Additionally, we assume that carbon production is proportionate to shoot leaf area, ignoring potential self-shading in larger growing plants. Combined this enables us to write the following equation for the growth dynamics in the root system:

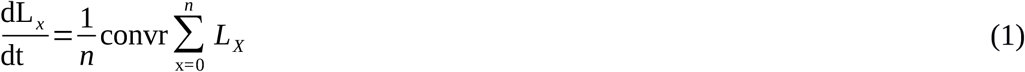

where *x* is the index indicating the number of the root system compartment modeled and *n* is the total number of root compartments considered, with-unless specified otherwise-all root compartments obtaining an equal fraction 1/*n* of the total energy available for root growth. Additionally, conv represents the convergence of shoot area (assumed proportionate to overall root system length) via carbon production into root length increase, and *r* represents a growth rate function that we will later use to incorporate the dependence of growth rate on external and internal nitrate levels. Note that this equation will result in exponential root growth dynamics, consistent with the growth dynamics observed for young *Arabidopsis thaliana* plants in absence of resource limitations, competition or stress (Guan et al., 2014).

To incorporate the effects of both external, environmental as well as internal, plant systemic nitrate status on growth, we need to incorporate the uptake of external nitrate and the subsequent intra plant nitrate dynamics in our model. For this we incorporate both the dynamics of local, internal nitrate levels (*N*_*i*_) that we model per root system compartment, as well as an overall, systemic nitrate level *N*_*s*_. Internal local nitrate levels increase through uptake of external nitrate (*N*_*e*_) from the environment and loose nitrate through transporting it to the system level nitrate pool. This nitrate pool in turn loses nitrate to turnover and maintenance of plant tissue as well as exudation. To take into account the uptake of external nitrate by both high and low affinity transporters (Crawford and Glass, 1998), we consider both saturated and non-saturated nitrate uptake. Combined this leads to the following equations:

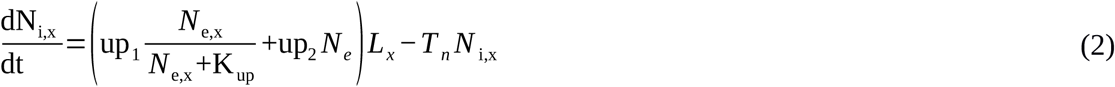

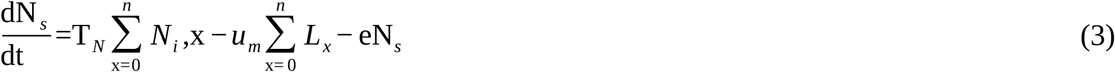

where up_1_ is the maximum uptake rate of the high affinity transporters, *K*_up_ is the concentration at which these high affinity transporter operate at half maximum velocity, up_2_is the uptake rate of low affinity transporters that for simplicity are assumed not to saturate, *T*_*N*_ is the rate of transport of nitrate from the local to the systemic nitrate pool, *u*_*m*_ is the rate of nitrate loss to tissue maintenance and turnover and *e* is the rate of nitrate loss to exudation. Note that changes in external nitrate levels are ignored in our model.

#### 2.1.2 Single root, split root and patch experiments

Varying the number of root compartments we simulate and exposing these compartments to different environmental nitrate levels enables us to simulate various experimental setups. To investigate the effect of changes in overall, homogeneous nitrate concentration on root system growth, we can simply apply a single root compartment. If instead we aim to simulate split root experiments in which different root halves (potentially) experience different external nitrate levels, we can apply two root compartments. Finally, to investigate the role of isolated nitrate patches on root system growth, we can simulate a large number (in this study n=16) of root system compartments, with only one of these containing high nitrate levels. It is important to note that in absence of additional regulation, differences in the external nitrate levels to which different simulated root compartments are exposed will only result in different internal nitrate levels but not in differences in the compartment root system sizes (Eq. 1).

### 2.2 Dimensions and parametrization of the model

#### 2.2.1 Units of model variables

Note that since in our simplified model we work only with length, and not radius, volume or weight of the root system, for convenience concentrations are computed per unit length. Thus, *L*, length of (part of the) root system is in mm, *N*_*i*_, internal nitrate amount, and *N*_*s*_, systemic nitrate signalling amount, is in micromole, whereas 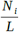 internal nitrate concentration and 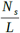 systemic nitrate signalling concentration is in micromole/mm.

#### 2.2.2 Parameter values

For the maximum uptake rate of high affinity transporters, depending on the study and the specific nitrate transporter studied, values between 0.3 and 8 micromole/g freshweight/h are reported (Crawford and Glass, 1998). Since we only incorporated a single, generalised high affinity nitrate uptake transporter in our baseline model, we choose an intermediate value of 3.6 micromole/g freshweight/h. Next, since in our simplified model all is per unit length, we need to convert this value to micromol/mm/h. For this we used data from a study on *Brachypodium* (Sasse et al., 2019), reporting an approximately 1.2:1 ratio between 1 g freshweight and 1 cm root length, resulting in a rounded off 0.6 micromole/mm/h value for up_1_. Similarly, for *K*_up_, also depending on study and nitrate transporter studied, values ranging from 6 to 100 micromole/L have been reported (Crawford and Glass, 1998), and we again take an intermediate value of 50 micromole/L.

Given the simplified nature of our model, for other parameters values could not be directly derived from available data. E.g. *T* _up_, the rate of transport of internal nitrate to the systemic nitrate pool, is in fact a compendium of the characteristics of nitrate transporters in phloem and xylem, as well as their numbers and distribution, the distance covered, etc. Therefore, instead we fitted *T* _up_ and up_2_ to reproduce the experimentally observed dependence of internal nitrate concentrations on external nitrate concentrations. For this we used data from a study by (Gruber et al., 2013), in which *Arabidopsis* plants were exposed to a range of external nitrate concentrations ranging from 110 to 11000 micromole/L. Internal nitrate concentrations reported in (Gruber et al., 2013) are shoot/root levels of 72/48; 66/45, 48/39 and 29/na mg/g dryweight for externally applied nitrate of 11400, 550, 275 and 110 micromole/L respectively. First, the not available (na) root internal nitrate level for 110 micromole/L external nitrate was extrapolated from the available root and shoot values, assuming that for 110micromole/L the ratio between shoot and root nitrate levels is 1.1, resulting in a value of 26 mg/g dryweight. Thus, shoot nitrate levels vary 2.5 fold, and root nitrate levels vary 1.8 fold over a 100 fold change in external nitrate levels. Next, we converted the internal nitrate levels to micromole/g freshweight by assuming a 4 fold weight difference between fresh and dry weight (effectively assuming 75% of plant mass consists of water), consistent with classical experimental values (van de Sande-Bakhuyzen, 1928), and by using the molecular volume for nitrate of 62,0049 g/ mol. This results in converted root internal nitrate levels of 0.194, 0.181, 0.157 and 0.105 micromol/g freshweight. Finally, since we describe root growth in terms of length increase, we converted these internal nitrate levels to micromole/mm. For this we again used data from a study on *Brachypodium* reporting both overall root length and root system weight values, from which we derived a 1.2:1 ratio between 1 g freshweight and 1 cm root length (Sasse et al., 2019). After this final conversion we obtained root internal nitrate levels of 0.161, 0.151, 0.131 and 0.087 for 11400, 550, 275 and 110 micromole/L external nitrate respectively. By taking *T*_up_=3.8 and up_2_=0.000006 we obtained a good fit for the dependence of internal nitrate levels on external nitrate between the model and the experimental data (see Fig 2A in Results section, compare black line with green circles).

**Figure 2.**
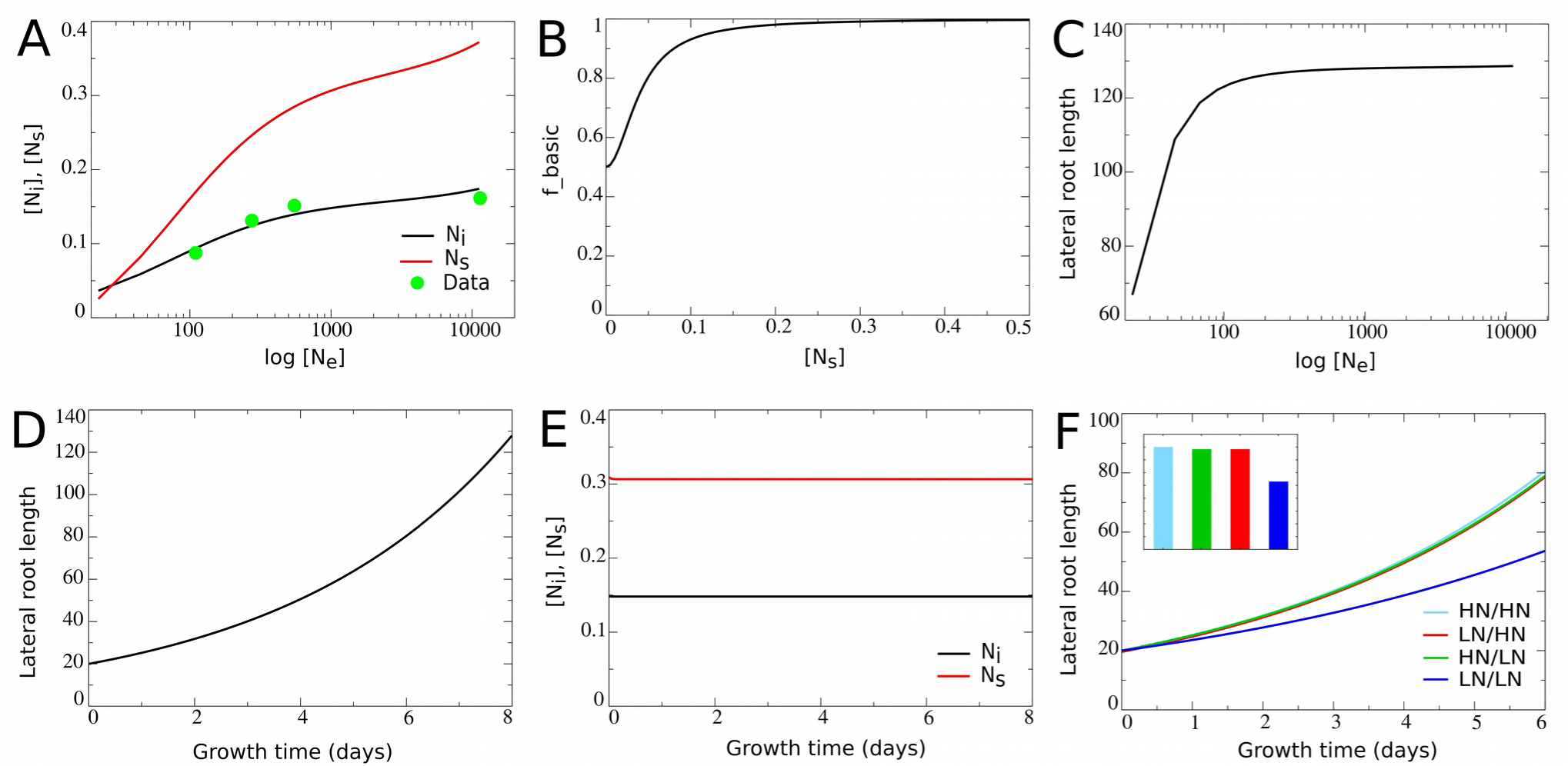
Incorporating a saturating dependence on systemic nitrate levels. **(A)** Internal and systemic nitrate levels as a function of external nitrate for the basic model settings in which growth is nitrate independent. **(B)** Model survival response: growth rate decrease as a function of systemic nitrate levels. **(C)** Cumulative root system length after 8 days of growth (in mm) as a function of external nitrate levels in the model including the survival response. **(D)** Exponential root growth dynamics for an external nitrate level of 1000 mM. **(E)** Internal and systemic nitrate dynamics for the root growth shown in D. **(F)** Growth dynamics of individual root halves in split root experiments. For plants experiencing identical nitrate concentrations in left and right halves growth dynamics of only a single root half are shown.

Parameter values for *u*_*m*_ (0.1) and *e* (1.5) were chosen such that systemic nitrate levels show a larger range of variation as a function of external nitrate compared to the local internal nitrate levels (Fig 2A, compare black and red lines). The reason for doing this is that systemic nitrate level is known to affect root system growth in various ways, at different systemic nitrate levels. A survival response, during which root growth is strongly repressed occurs for very low systemic nitrate levels. In contrast, for somewhat higher systemic nitrate levels a foraging response promoting root growth is induced. Finally, for very high systemic nitrate levels, systemic repression reduces root growth (Giehl and Wirén, 2014). In order to incorporate these different effects into our model in a robust manner, we should be able to activate these effects at sufficiently different systemic nitrate concentrations, requiring a large enough range of systemic nitrate levels to occur in our model. As a final parameter we need to determine the value for conv. For this, we make use of the fact that we can write an analytical solution for Eq.1:

*L* (*t*) =L(0)*e*^convrt^

Typically, in split root experiments, split root conditions are started when the first order laterals have grown to a size of 2-4 cm (see e.g. (Guan et al., 2014)). We therefore take the start size of a single side of the root system (with only one side present in case of an unsplit root system) as *L* (0)=20 mm. Experimental data also indicate that the maximum cumulative length of secondary laterals in one root half after 10 days of treatment lies around 24 cm, and far exceeds the primary lateral root length (Guan et al., 2014). Thus, we assume that after 10 days the maximum size of half the root system is 36 cm (1.5 times 25 cm) and hence the maximum size for the whole root system size lies around 72 cm, so *L* (240 *h*)=720 mm. Since this is the maximum root system size occurring under favorable conditions, i.e. conditions promoting growth, we assume that r= 1.5 rather than being equal to 1. Together this gives:

720=20 *e*^conv 1.5240^

from which we solve conv= 0.01.

Combined this results in the parameter settings shown in Table 1.

**Table 1.** Units and values of model parameters.

| Model parameters | Units | Values |
| --- | --- | --- |
| $up_1$ | micromole/mm/h | 0.6 |
| $K_{up}$ | micromole/L | 75 |
| $up_2$ | micromole/mm/h | 0.000006 |
| $N_e$ | micromole/L | 110-11000 |
| $T_{up}$ | /h | 3.8 |
| $u_m$ | micromole/mm/h | 0.1 |
| $e$ | /h | 1.5 |
| conv | /h | 0.001 |

### 2.3 Model extensions

#### 2.3.1 Local and systemic CEP dynamics

To model the effect of nitrate demand signalling on preferential root foraging (see Results section), we extended our model with local nitrate dependent production of CEP, which subsequently is transported to a systemic CEP pool, in which it turns over with a certain rate. To describe these dynamics, we extended our model with the following equations:

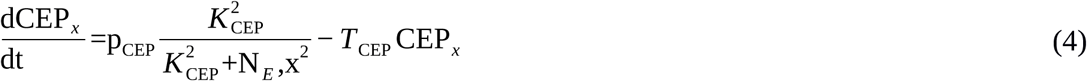

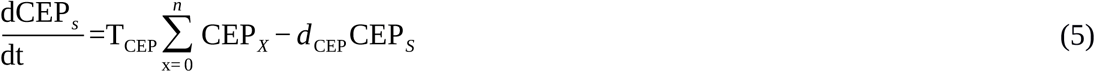

where *p*_CEP_ is the maximum rate of CEP production, *K*_CEP_ is the external nitrate concentration at which CEP production reaches its half maximum rate, *T*_CEP_ is the rate of transport from the local to systemic CEP pool, and *d*_CEP_ is the degradation rate of CEP.

#### 2.3.2 Local and systemic CK dynamics

To incorporate the effect of nitrate supply signalling on preferential root foraging (see Results section), we added to our model local CK production, it’s transport to a system level CK pool, and it’s decay in that pool, using the following equations:

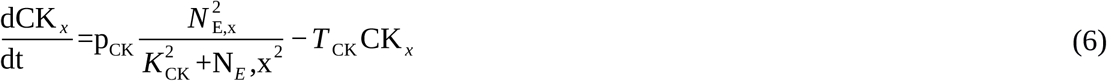

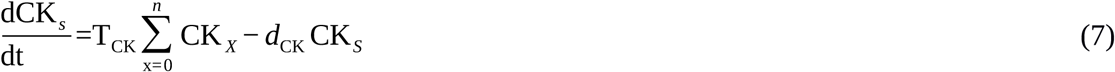

where *p*_CK_ is the maximum rate of CK production, *K*_CK_ is the external nitrate concentration at which CK production reaches its half maximum rate, *T*_CK_ is the rate of transport from the local to systemic CK pool, and *d*_CK_ is the degradation rate of CK.

#### 2.3.3 Parameter settings for model extensions

In the Results section, as well as above, we describe how our baseline model is extended to incorporate the various known aspects of internal and external nitrate status dependent growth regulation. Parameters, values and dimensions, involved in these model extensions are listed below, in Table 2.

**Table 2.**
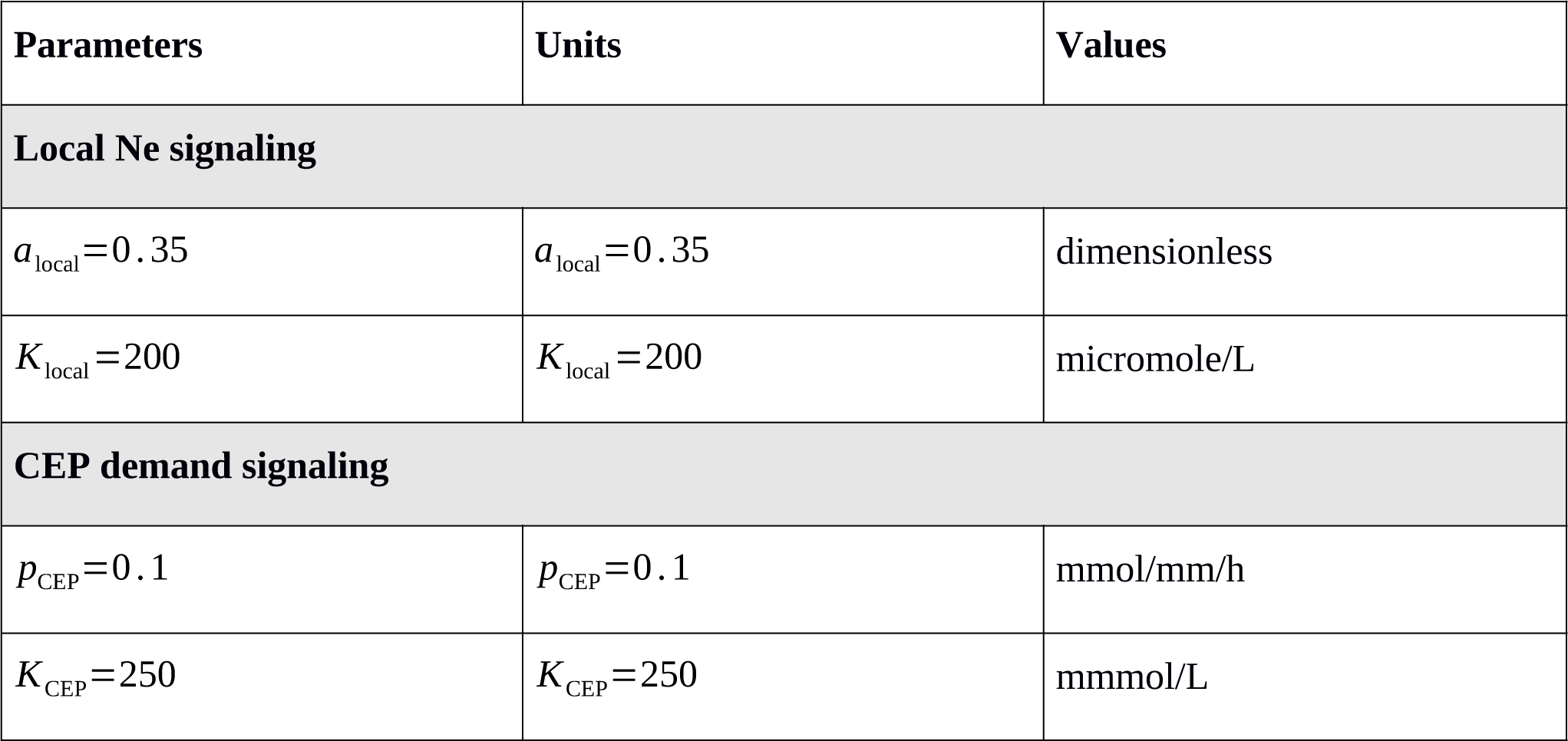
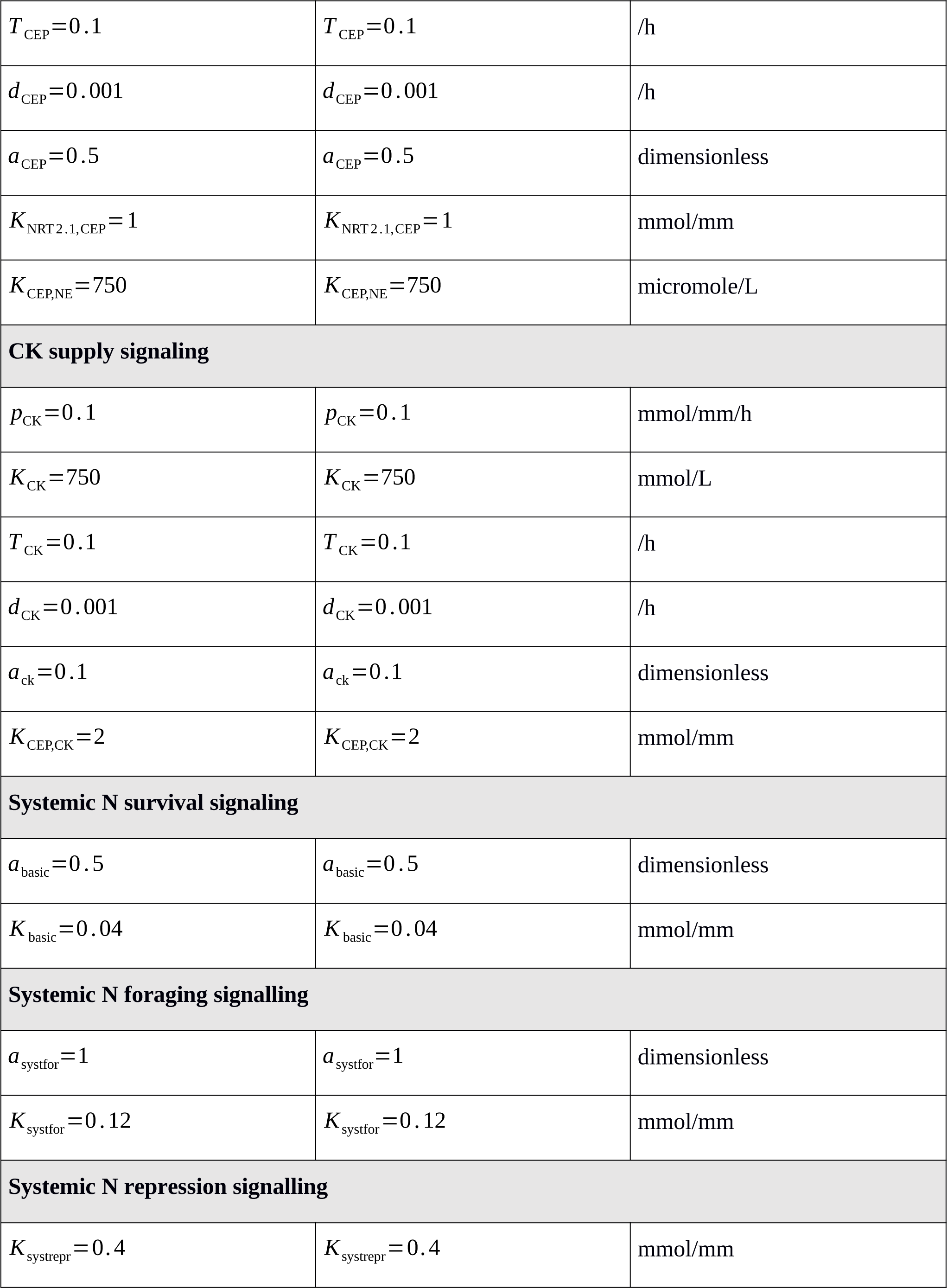
Parameter settings for model extensions.

| Parameters | Units | Values |
| --- | --- | --- |
| **Local Ne signaling** |  |  |
| $a_{local}=0.35$ | $a_{local}=0.35$ | dimensionless |
| $K_{local}=200$ | $K_{local}=200$ | micromole/L |
| **CEP demand signaling** |  |  |
| $p_{CEP}=0.1$ | $p_{CEP}=0.1$ | mmol/mm/h |
| $K_{CEP}=250$ | $K_{CEP}=250$ | mmmol/L |
| $T_{\text{CEP}}=0.1$ | $T_{\text{CEP}}=0.1$ | /h |
| $d_{\text{CEP}}=0.001$ | $d_{\text{CEP}}=0.001$ | /h |
| $a_{\text{CEP}}=0.5$ | $a_{\text{CEP}}=0.5$ | dimensionless |
| $K_{\text{NRT2.1,CEP}}=1$ | $K_{\text{NRT2.1,CEP}}=1$ | mmol/mm |
| $K_{\text{CEP,NE}}=750$ | $K_{\text{CEP,NE}}=750$ | micromole/L |
| **CK supply signaling** |  |  |
| $p_{\text{CK}}=0.1$ | $p_{\text{CK}}=0.1$ | mmol/mm/h |
| $K_{\text{CK}}=750$ | $K_{\text{CK}}=750$ | mmol/L |
| $T_{\text{CK}}=0.1$ | $T_{\text{CK}}=0.1$ | /h |
| $d_{\text{CK}}=0.001$ | $d_{\text{CK}}=0.001$ | /h |
| $a_{\text{ck}}=0.1$ | $a_{\text{ck}}=0.1$ | dimensionless |
| $K_{\text{CEP,CK}}=2$ | $K_{\text{CEP,CK}}=2$ | mmol/mm |
| **Systemic N survival signaling** |  |  |
| $a_{\text{basic}}=0.5$ | $a_{\text{basic}}=0.5$ | dimensionless |
| $K_{\text{basic}}=0.04$ | $K_{\text{basic}}=0.04$ | mmol/mm |
| **Systemic N foraging signalling** |  |  |
| $a_{\text{systfor}}=1$ | $a_{\text{systfor}}=1$ | dimensionless |
| $K_{\text{systfor}}=0.12$ | $K_{\text{systfor}}=0.12$ | mmol/mm |
| **Systemic N repression signalling** |  |  |
| $K_{\text{systrepr}}=0.4$ | $K_{\text{systrepr}}=0.4$ | mmol/mm |

## 3 Results

### 3.1 Establishing a baseline root growth model

To establish a baseline model for *Arabidopsis thaliana* root growth in which subsequent extensions can be built, we start with a single, non-split root that takes up external nitrate from the environment, transports this nitrate internally hereby resulting in the establishment of a systemic nitrate level, and grows (see Materials and Methods, Eqs 1-3).

In nature, soil nitrate levels have been found to vary 5 to 7 fold with depth in the soil (Angle et al., 1993; Jin et al., 2015), 3-5 fold with seasonal changes (Taylor et al., 1982; Weil and Brady, 1996; Hellebrand et al., 2005), and 3 fold even between similar soils (Jin et al., 2015). Combined, this suggests that plant roots experience variations in soil nitrate levels of up to two orders of magnitude. At the same time experimental data how that 100 fold changes in external nitrate results in only 1.82 to 2.5 fold changes in rootleaf and root nitrate levels (Gruber et al., 2013). To be able to simulate this we fitted model parameters to experimental data (see Materials and Methods section), initially assuming no nitratees dependent growth regulation (r= 1). Figure 2A plots simulated local internal and systemic nitrate levels as a function of external nitrate and shows a good agreement between simulated and experimentally obtained internal nitrate levels.

Next, we introduce the first, basic dependence of root growth on nitrate levels. Obviously, plant growth ultimately depends on the carbon generated through photosynthesis, which in turn is highly dependent on the protein Rubisco and as such also dependent on nitrate levels. Additionally, plants have been shown to display a survival response for low external (and hence systemic) nitrate levels, repressing root growth via the CLE-CLV1 module (Araya et al., 2014) as well as through ACR4/ AXR5 (Giehl et al., 2014). Based on this we incorporate a saturating dependence of growth rate on systemic nitrate levels writing:

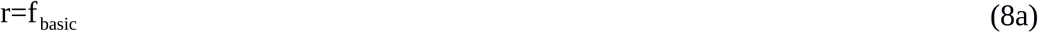

with

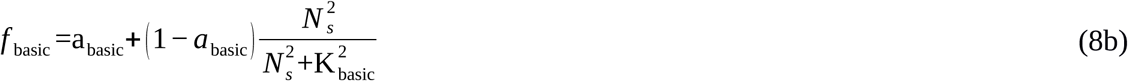

where *a*_basic_ is the *N*_*s*_ dependent and (1 *− a*_basc_) the *N*_*s*_ independent fraction of *f* _basic_, and *K* _basic_ is the systemic nitrate level at which the systemic nitrate dependent fraction of the growth rate is half maximal. Based on the systemic nitrate levels occurring in Fig 2A, to ensure that survival responses occur only at very low nitrate levels, we choose *K* _basic_=1. Next we need to decide on the value for *a*_basic_. In split root experiments in which both root halves are exposed to low or even absent nitrate, some root growth still occurs (Ruffel et al., 2011; Guan et al., 2014; Mounier et al., 2014). This likely results from the exposure to external nitrate prior to the start of the split root experiments causing the presence of stored nitrate. Since we do not include nitrate stores in our simple model, yet aim to simulate split root experiments, we model this limited reduction of root growth by the survival response by using a value of a= 0.5. Combined this results in the dependence of *f* _basic_ on *N*_*s*_ as shown in Fig 2B. Figure 2C shows root system length after 8 days of growth as a function of external nitrate levels based on these growth rate settings, showing a saturating effect of external nitrate on overall root system size yet without root growth being fully abolished at very low external nitrate levels. When plotting the temporal dynamics of root growth at a single external nitrate level (*N*_*e*_=1000), we obtain an exponential increase in root growth length over time (Fig 2D), as has been observed experimentally (Guan et al., 2014). Additionally, we see that after an initial transient, steady state local and systemic nitrate concentrations is reached despite continued growth (Fig 2E).

Next we investigate root growth dynamics produced when simulating classical split root experiments, in which root halves are exposed to either very low (25 mM) or high (5000 mM) external nitrate levels. Note that for clarity, for the situations in which both root halves are exposed to the same nitrate level only the length of a single root half is shown. It can be seen that root growth is less when both root halves are exposed to very low nitrate levels, yet does not differ much between the situation when only one or both root halves are exposed to a high nitrate level (Fig 2F). This is consistent with the applied saturating dependence of root growth on systemic nitrate levels. Additionally, we see that no differences between left and right root halves of plants experiencing heterogeneous external nitrate conditions occur. This logically follows from the fact that growth in the current model settings only depends on systemic but not local internal nitrate levels.

### 3.2 Local nitrate signalling

Local nitrate levels have been shown to affect lateral root growth root. One of the key players involved in this local response is the nitrate transceptor NRT1.1. For low external nitrate levels, NRT1.1 has been shown to function as an auxin importer, resulting in the reduction of local auxin levels and thereby inhibiting lateral root growth. In contrast, for higher external nitrate levels, NRT1.1 does not transport auxin, therefore not having this negative effect on root growth (Krouk et al., 2010). Additionally, for higher external nitrate levels NRT1.1 positively influences auxin signalling and hence lateral root growth via the AFB3, NAC4, OBP4 pathway (Vidal et al., 2010, 2013, 2014), as well as ANR1 (Remans et al., 2006).

To investigate the contribution of local nitrate sensing to preferential foraging with our model we extended our baseline model by incorporating a root growth promoting function that depends in a saturating manner on the local external nitrate level:

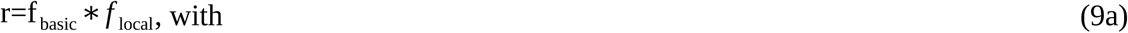

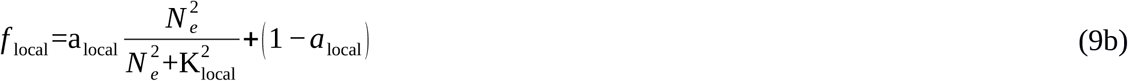

,where *a*_local_ represents the *N*_*e*_ dependent and (1 *− a*_local_) the *N*_*e*_ independent fraction of *f* _local_, and *K* _local_ represents the external nitrate concentration at which the external nitrate dependent fraction reaches half of its maximum value.

Parameter values are chosen such (see Table 2) that a baseline growth rate of 0.65 arises if no external nitrate is present, with growth rates increasing to 1 as external nitrate increases (Fig 3A). As a consequence, relative to an external nitrate level of say 250 mM which results in a growth rate of approximately 0.8, both external nitrate dependent decreases and increases in growth rate may occur, consistent with the above described experimental data. In Figure 3B, outcomes of split root simulations under these new model settings are shown. We see that with the impact of external nitrate levels added to that of systemic nitrate levels, root length differences between plants experiencing only very low or only very high nitrate concentrations have increased (Fig 3B, dark blue versus light blue). Furthermore, as expected, we now see that in plants experiencing heterogeneous external nitrate concentrations, an asymmetry in the length of the two root halves occurs with the root half experiencing higher nitrate levels growing longer. However, root system length on the high nitrate side is lower compared to the situation in which both root halves experience high nitrate levels. On a similar note, root system length on the low nitrate side are higher as compared to the situation in which both root halves experience low nitrate levels, the reverse of what is observed experimentally (Fig 3B). This can be easily understood from the dependence of root growth on both*r*, and hence local nitrate, as well as on overall root system size*L*(assumed proportional to shoot size and hence carbon availability for growth) in our model. In case of the heterogeneous split root system, growth is stimulated in one root half, resulting in less root length increase as compared to when two halves and more root length increase as compared to when none of the halves are stimulated by local nitrate.

**Figure 3.**
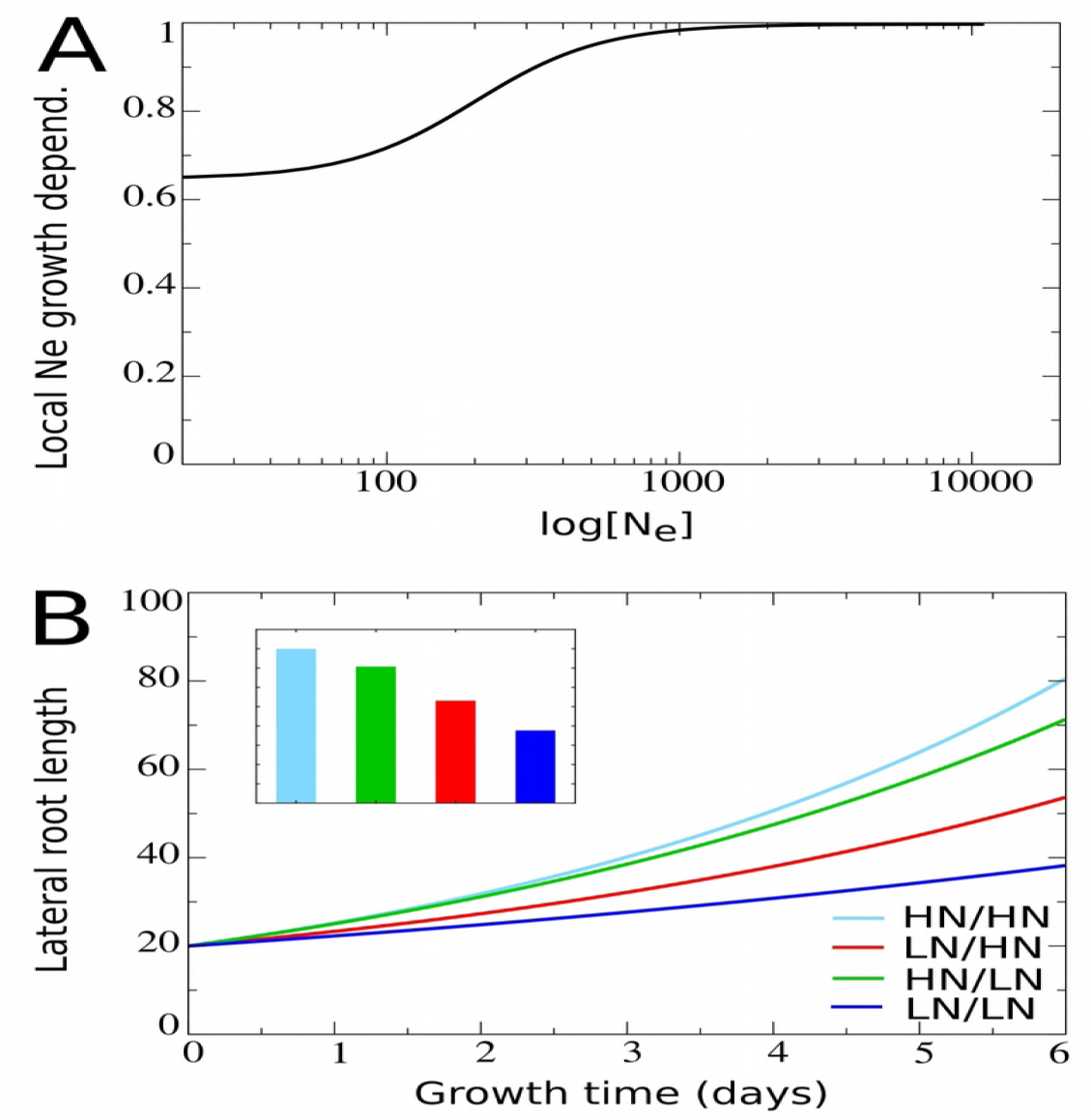
Including the dependence of root growth on local nitrate levels. **(A)** Model local stimulation response: growth rate dependence on local external nitrate levels. **(B)** Growth dynamics of individual root halves in split root experiments. Colors same as in Fig 2F.

### 3.3 Systemic demand signalling

In contrast to the results obtained above, plants show a preferential increase in lateral root lengths at the high nitrate side as compared to plants experiencing high nitrate at both sides (Fig 1B), suggesting the involvement of a growth promoting systemic demand signal. Recently, at least part of such a systemic nitrate lack signalling system has been uncovered. It was shown that under low external nitrate levels lateral roots produce CEP peptides, which in the shoot bind to CEP receptor and cause the production of CEP downstream signals that travel back to the root. CEP downstream signals combined with the local presence of sufficient nitrate subsequently leads to the upregulation of NRT2.1 (Tabata et al., 2014; Ohkubo et al., 2017), and results in upregulation of nitrate uptake as well as nitrate dependent root growth (Little et al., 2005; Remans et al., 2006; Naz et al., 2019) (Fig 4A).

**Figure 4.**
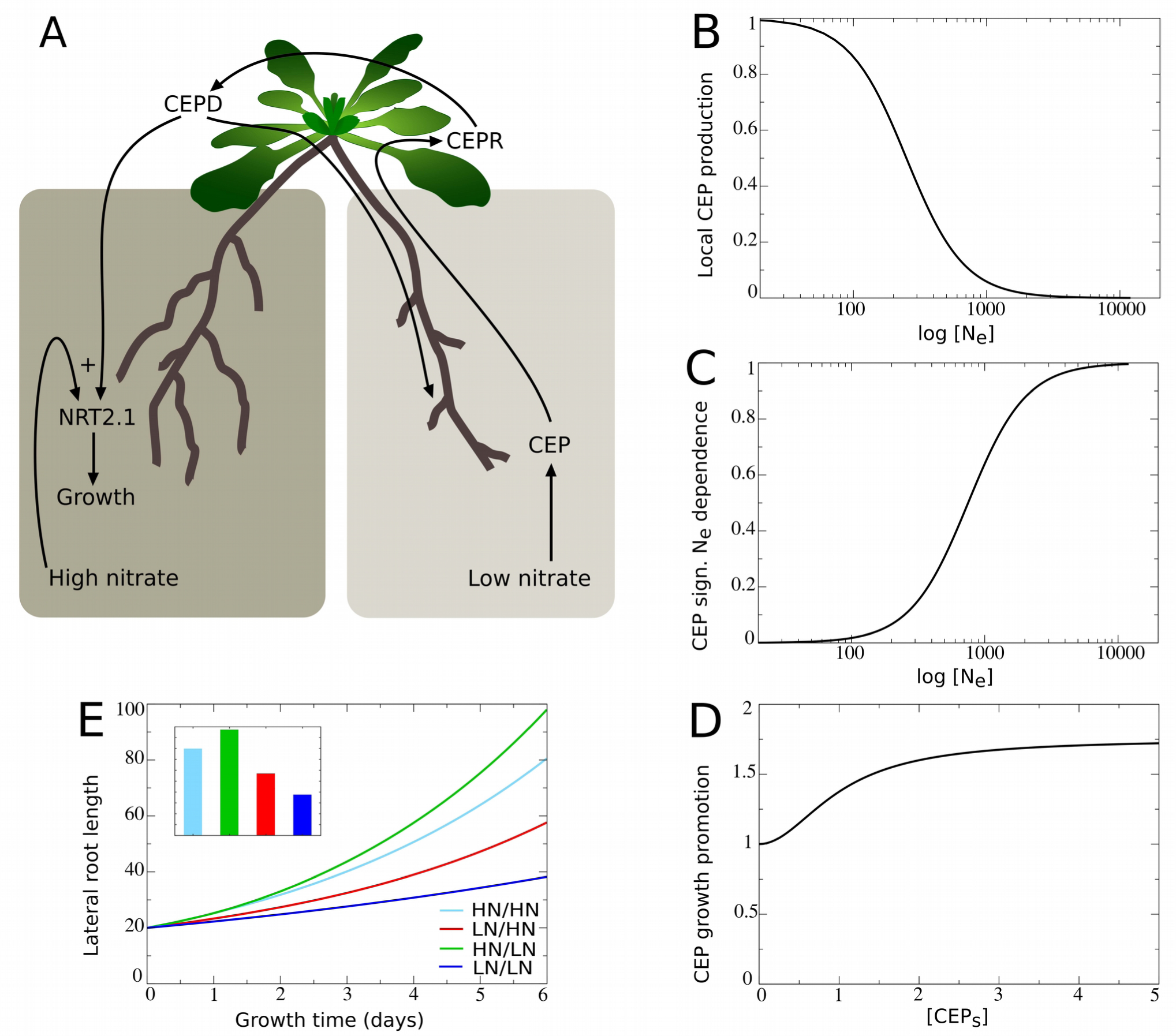
Including the dependence of root growth on CEP-mediated demand signalling. **(A)** Schematic depiction of the CEP-mediated demand signaling system, showing the low nitrate induced production of CEP (1), the CEP and local nitrate dependent upregulation of NRT2.1 (2), and the NRT2.1 dependent stimulation of root growth (3). **(B)** Dependence of the rate of CEP production on the low nitrate side on local external nitrate levels. **(C)** Modulation of CEP signalling effect on root growth promotion on the high nitrate side (D) on local external nitrate. **(D)** Dependence of root growth promotion on the high nitrate on systemic CEP levels. **(E)** Growth dynamics of individual root halves in split root experiments. Colors the same as in Fig 2F.

To incorporate this mechanism in our model we add the local nitrate dependent production of CEP, with CEP production decreasing in a saturating manner with increasing external nitrate levels (Fig 4B). Additionally, we model the transport of CEP to the shoot, where it enters the systemic CEP pool and has a certain rate of turnover (See Materials and Methods, Eq 4, 5).

To restrict the number of variables included in our model, rather than explicitly incorporating CEP receptors and downstream signals, we incorporated a direct dependence of the promotion of root growth by systemic CEP signaling levels (CEP_*S*_). Since CEP mediated signaling has been shown to promote, yet not repress root growth, our function is chosen such that in absence of systemic CEP and other regulations, a baseline growth rate of 1 occurs, while in presence of systemic CEP this growth rate is enhanced (Fig 4D). We thus now model growth rate as:

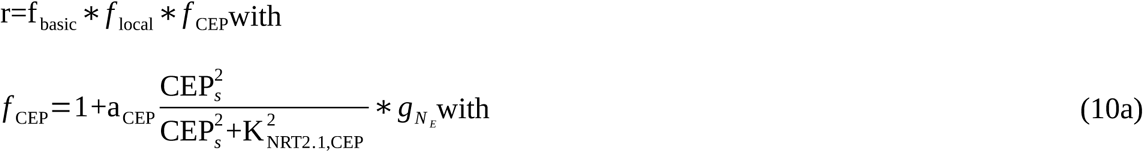

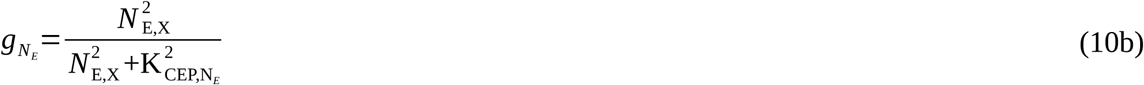

where *a*_*CEP*_ is the maximum CEP signalling induced increase in growth rate, *K* _NRT2.1, *CEP*_ is the CEP level at which this growth rate increase reaches it’s half maximum value, and *K_CEP, N_* is the local nitrate level at which this growth rate increase reaches it’s half maximum value. By multiplying the CEP dependent part of this function with *g*, which depends in a saturating manner on local external nitrate (Fig 4C) we incorporate that CEP mediated growth promotion only occurs in the presence of local nitrate.

Figure 4E shows how combining this mechanism with the earlier systemic and local nitrate dependent growth regulating mechanisms results in the preferential enhancement of root growth at the high nitrate side in split root plants experiencing heterogeneous nitrate conditions.

### 3.4 Systemic repression

In addition to the high nitrate side having longer lateral roots in heterogeneous as compared to homogeneous conditions, root length under homogeneous high external nitrate conditions has been observed to be very low (Fig 1B). This reduction of overall lateral root length under high nitrate has been attributed to systemic repression (Fig 1A) (Giehl and Wirén, 2014). It is considered as an adaptive response, resulting in the reduced investment of a plant in its root system under conditions of superfluous nutrient availability, and is mediated through the repression of auxin sensing via the AFB3, NAC4, OBP4 pathway (Vidal et al., 2010), and may also involve the HNI9 mediated repression of nitrate transport (Girin et al., 2010).

To further improve the realism of our model we therefore incorporated a function describing the decrease of root growth rate with systemic nitrate levels:

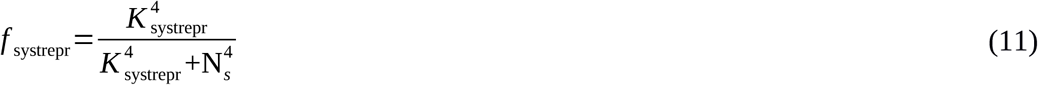

where *K*_systrepr_ is the systemic nitrate level at which *f* _systrepr_ has decreased to half its maximum value. Based on the range of systemic nitrate levels observed in Figure 2A we choose *K*_systrepr_=10, ensuring repression only occurs for very high internal nitrate levels (Fig 5A). As expected, when comparing Figure 5B to Figure 4E mostly the root lengths of the plant experiencing high nitrate levels on both sides have decreased.

**Figure 5.**
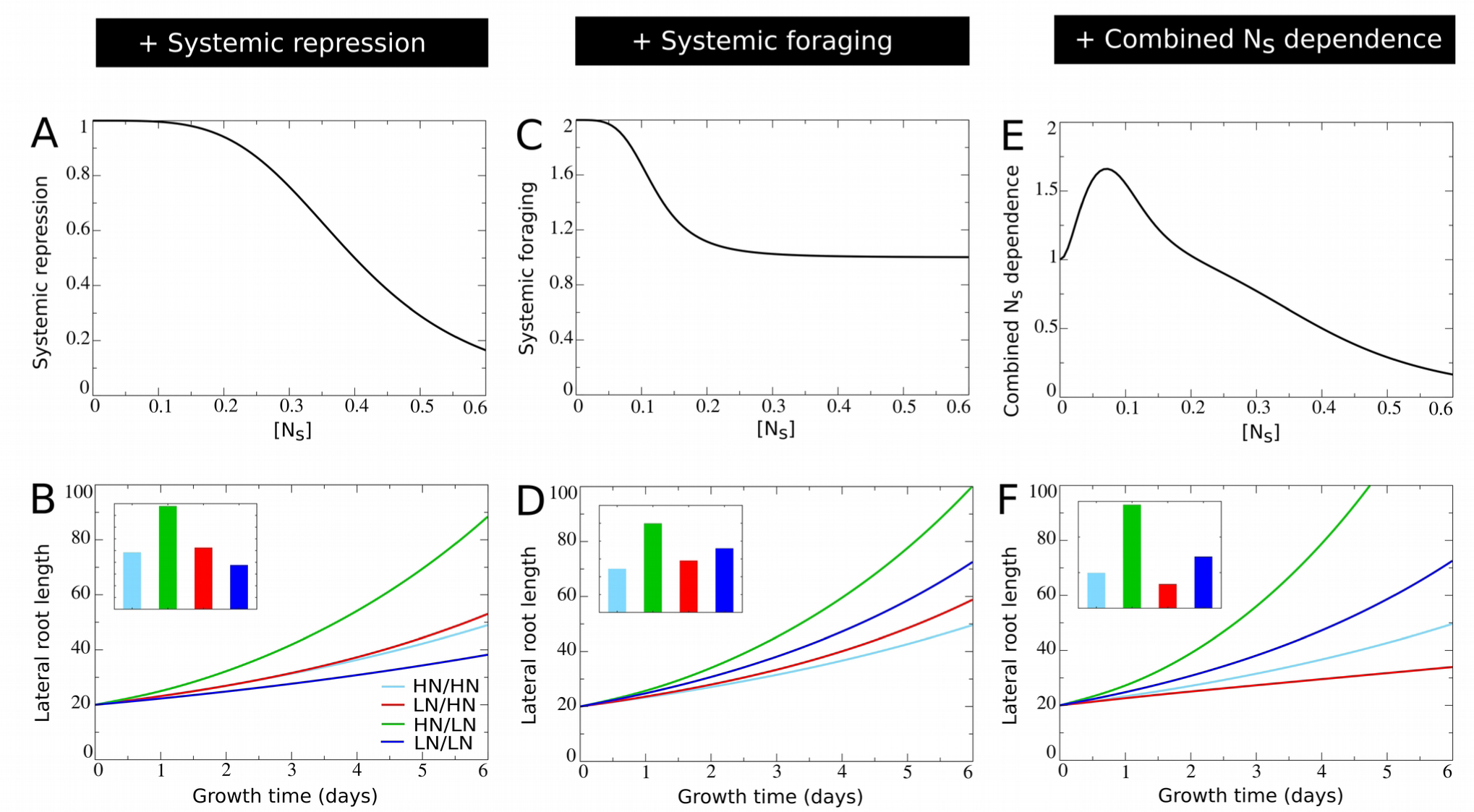
Incorporating systemic repression, systemic foraging and competition for carbon. **(A)** Model systemic repression response: growth rate decrease as a function of increasing systemic nitrate levels. **(B)** Growth dynamics of individual root halves in split root experiments when a systemic repression response is added. **(C)** Model systemic foraging response: growth rate increase as a function of decreasing systemic nitrate levels. **(D)** Growth dynamics of individual root halves in split root experiments when a systemic root foraging response is added. **(E)** Growth rate modulation occurring as a combination systemic survival, foraging and repression responses (shown is *f* _basic_ ∗ *f* _systrepr_ ∗ *f* _systfor_). **(F)** Growth dynamics of individual root halves in split root experiments when competition for carbon is added. For B, D and F colors are same as in Fig 2E.

### 3.5 Systemic foraging

Thusfar our model offers an explanation for only one half of the preferential root foraging phenotype. *In planta*, in addition to a preferential increase in lateral root lengths on the high nitrate side, also a preferential decrease on the low nitrate side is observed. That is, lateral root length on the low nitrate side is lower than the lateral root lengths of plants experiencing low nitrate levels on both sides. In our current model settings, this is difficult to reproduce due to the very low root lengths occurring for plants experiencing homogeneous low nitrate levels. However, in plants, in between the extremely low systemic nitrate levels inducing a survival response and the very high nitrate levels inducing systemic repression, a third growth response occurs. This response is referred to as a root foraging response, and occurs for intermediate low systemic nitrate levels (Giehl and Wirén, 2014). Since the foraging response promotes growth at low systemic nitrate levels, yet does not repress growth at high nitrate levels, our function is choosen such that for higher systemic nitrate level a baseline growth rate of 1 occurs, while in presence of low systemic nitrate levels this growth rate is enhanced (Fig 5C). Again, we incorporated this growth affecting mechanism incrementally to our model using the following equation:

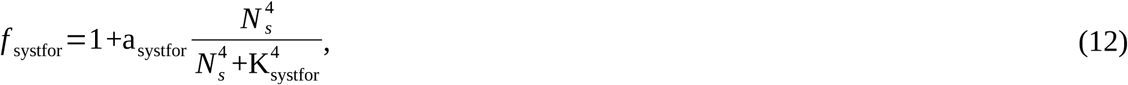

with *a*_systfor_ the amplitude with which low *N*_*S*_ stimulates growth, which becomes half maximal at an *N*_*S*_ concentration of *K*_systfor_. We choose *K*_systfor_=3, ensuring it to occur for higher *N*_*S*_ levels than the survival response (*K* _basic_=1) and for lower *N*_*S*_ levels than the systemic repression response (*K*_systrepr_=10) (Fig 5C). In Figure 5E the combined effect of systemic nitrate on root growth rate, incorporating, survival, foraging and systemic repression responses is shown. Incorporating our foraging growth response leads to an elevation of root growth of plants experiencing homogeneous low nitrate, finally resulting in a lower root length at the low nitrate side of plants experiencing heterogeneous nitrate levels as compared to plants experiencing homogeneous low nitrate levels (Fig 5D).

### 3.6 Carbon allocation

While we have meanwhile incorporated all major known local and systemic nitrate effects on root growth we still observe that root length at the low side of a plant experiencing heterogeneous nitrate levels is longer than in a plant experiencing homogeneous high nitrate conditions (Fig 5D). This contrasts with available experimental data (Fig 1B) (Ruffel et al., 2011; Guan et al., 2014; Mounier et al., 2014). It is of course likely that the simplified nature of our model limits its abilities to reproduce all aspects of plant root nitrate foraging, and that certain important regulatory mechanisms remain to be discovered. Still, we reasoned that an important growth regulatory aspect is was missing in our model, namely carbon allocation. Besides being regulated by hormones, microRNAs, peptides and gene expression, plant organ growth foremost depends on available carbon. Thusfar, we assumed both root halves to get equal amounts of the carbon available for the root system (See Materials and Methods, Eq. 1). However, it is well known that the allocation of carbon to organs strongly depends on their sink strength and hence potential growth rate (Marcelis, 1996). Additionally, a correlation between nitrate presence, root growth and carbon allocation has been reported in soybean (Fujikake et al., 2003). To incorporate this aspect into our model we modify Eq. 1 into:

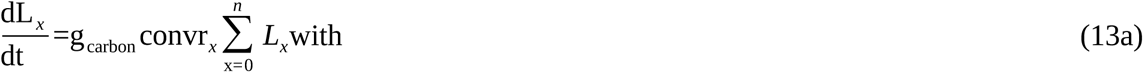

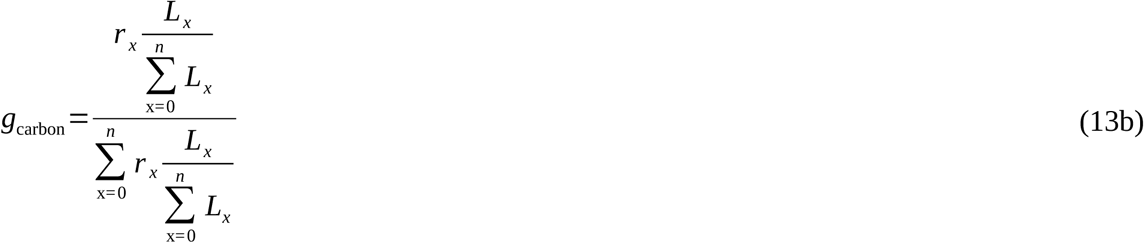

Put simply, the 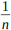 fraction in Eq. 1 that represented all *n* root compartments obtaining an equal fraction of carbon resources is replaced by a factor *g*_carbon_ describing carbon allocation as a function of relative growth rate (which would simply be 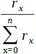), yet with the growth rates of the compartments weighted based on their relative size 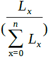.

Figure 5F shows how incorporating of this competition for carbon resources amplifies differences in root length between the high and low nitrate experiencing side of plants experiencing heterogeneous nitrate, and how this results in the low nitrate side now also having shorter root lengths than a plant exposed on two sides to high nitrate.

### 3.7 Role of systemic CK signalling

It has been recently demonstrated that systemic signalling occurring via root produced cytokinins also plays an important role in preferential root foraging under heterogeneous nitrate conditions. Based on the observation that triple ck biosynthesis mutants show hardly an increase in LR length on the high nitrate side compared to homogeneous nitrate conditions, this CK based signalling system was interpreted as a demand signal (Ruffel et al., 2011; Poitout et al., 2018). Still, root CK production correlates with local root nitrate levels, suggesting that it is more natural for CK to be a supply signal (Takei et al., 2002). This implies that for the demand driven upregulation of lateral root growth to function not only local nitrate presence signalling is required but also systemic supply signalling. Consistent with this interpretation is that the demand-signalling driven upregulation of NRT2.1 at the high nitrate side is largely abolished in the triple ck biosynthesis mutants (Poitout et al., 2018). Based on this we assume that nitrate dependent CK signalling influences the efficiency of CEP based signalling (Fig 6A). This could occur either through affecting CEP stability, CEP receptor numbers or affinity, or even TCP20 effectiveness (see (Guan et al., 2014)).

**Figure 6.**
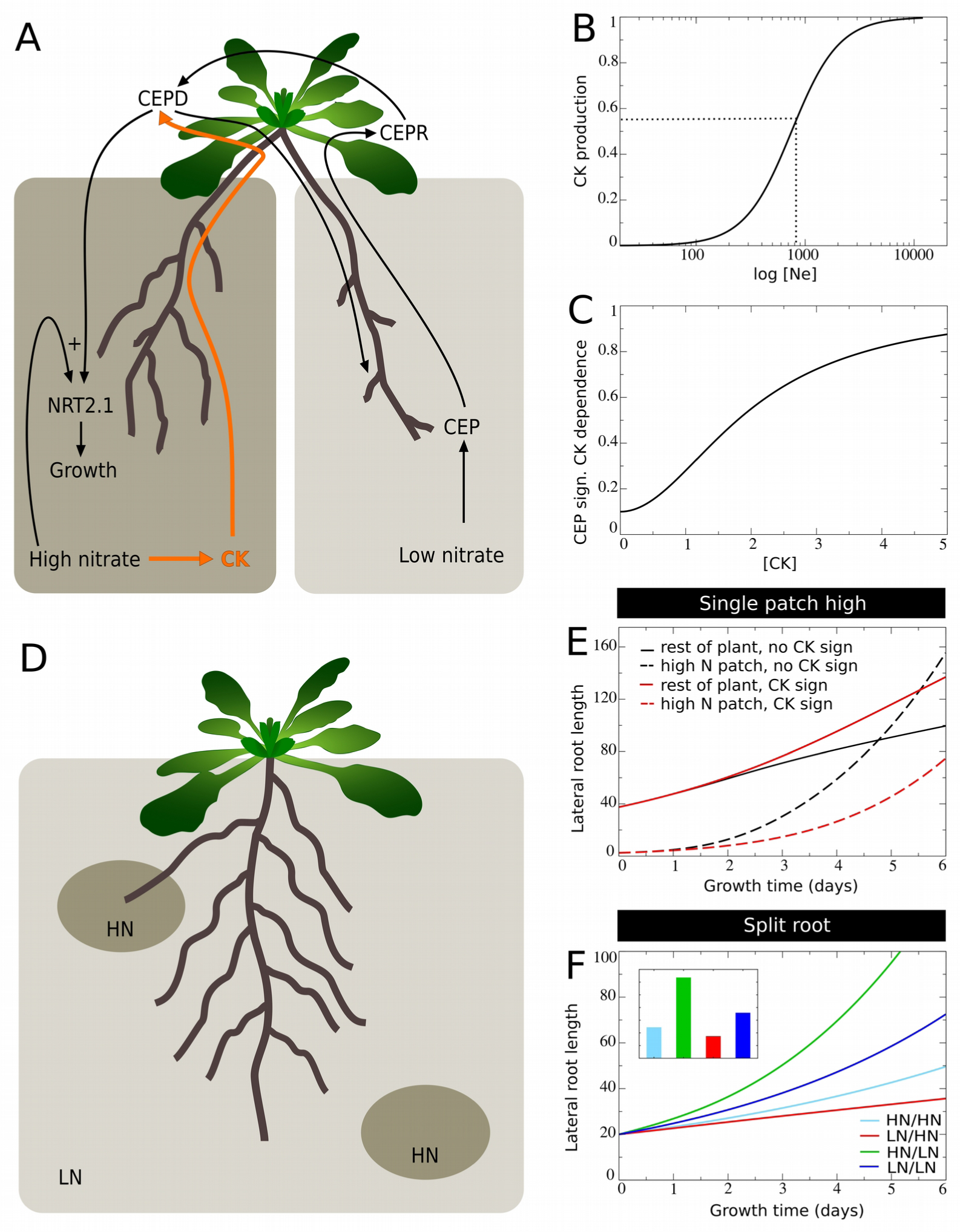
Incorporating systemic supply dependence of systemic demand signalling. **(A)** Schematic depiction of the proposed modulation of CEP demand signalling by CK supply signalling. **(B)** Production of CK as a function of external nitrate levels. **(C)** CK dependent modulation of CEP signalling strength. **(D)** Plant growing in a patchy nitrate environment, with part of its root system in a high nitrate patch and another high nitrate patch remaining to be discovered. **(E)** Growth dynamics in single compartment experiencing high nitrate and cumulative growth dynamics in other low nitrate experiencing compartments when CK supply signalling is not (black lines) or is (red lines) modulating CEP demand signalling strength. **(F)** Growth dynamics of individual root halves in split root experiments when CK supply signaling modulates CEP demand signalling. Colors same as in Fig 2E.

To incorporate this into our model, we simulate that local nitrate presence leads to local cytokinin production, which is subsequently transported shootward, resulting in a systemic cytokinin pool (CK_*s*_, Eq 6-7, Fig 6B). We subsequently redefine the CEP dependent growth control as:

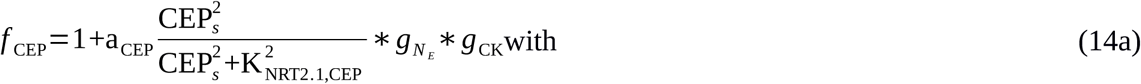

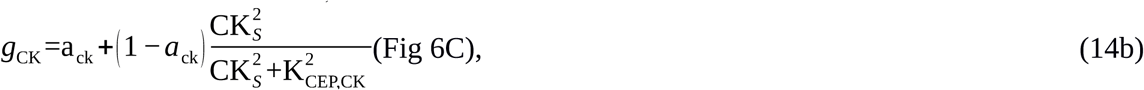

where *a*_*CK*_is the CK independent and 1 *− a_CK_* the CK dependent fraction of CEP signalling, and *K_CEP, CK_*is the CK level at which the CK dependent fraction of CEP signalling is at the half of its maximum level. causing the fraction1 *− a*_ck_of CEP dependent growth stimulation to depend on both local nitrate presence as well as systemic CK signalling.

An important question now is why plants, in addition to a “local nitrate presence signal” (*g*_*N*_), also would use a “systemic nitrate supply signal” (*g*_CK_). We hypothesize that while the local presence signal may serve to identify where to perform the preferential foraging growth, the systemic supply signal may serve to identify the extent of preferential foraging that is required by weighing supply and demand signals against one another. To investigate this, we simulated an “unbalanced” situation in which we partition the root system into n= 16 compartments, with only one of these compartments being exposed to high nitrate levels, and all others being exposed to low nitrate levels. This way, we simulate a root system exposed to a single high nitrate patch (Fig 6D).

In Figure 6E we plot the cumulative root length in the single compartment experiencing high nitrate level as well as the summed cumulative root length in all other compartments experiencing low nitrate levels. We do this for our previous model settings in which CEP signalling was CK independent (Eq. 10), and for our new model settings in which CEP signalling is CK dependent (Eq. 14). Initially, all root compartments have an equal size, causing the summed length of not high nitrate experiencing compartments to initially be 15 times higher. We see that for both situations, due to the high production of CEP in 15 out of 16 root compartments, root growth is strongly stimulated in the single high nitrate levels containing compartment. Still, when taking into account CK signalling, growth promotion in the nitrate patch is substantially less pronounced, and therefore growth reduction in the other compartments is also far less pronounced.

In Figure 6F we show a standard split root experiment, now with CEP signalling being CK dependent. Compared to Figure 5F results are highly similar. Thus, if, as is the case in a split root system in which one half experiences high and one half experiences low nitrate levels, CK supply and CEP demand signals are quantitatively balanced, the requirement for CK signalling hardly affects root growth. If instead demand by far outstrips supply, CK signalling prevents an excessive growth response at a single high nitrate patch, preventing the full collapse of root growth in other patches that may potentially reach other high nitrate patches (Fig 6D).

### 3.8 Mutants

Mutations in NRT1.1, CEP signalling and CK production have all been shown to cause a significant reduction in preferential root foraging (Ruffel et al., 2011; Mounier et al., 2014; Tabata et al., 2014; Ohkubo et al., 2017; Poitout et al., 2018). It was based on their individual large effects that we assumed that these players affect preferential root foraging in a synergistic rather than additive fashion, and let us to model their effects in a largely multiplicative manner. With all players in place we now test whether mutations in either 3 indeed strongly reduces preferential root foraging.

To simulate a mutation in NRT1.1, we put *f* _local_ to a constant intermediate value of 0.6, additionally, since NRT1.1 not only functions as a nitrate sensor but also a nitrate transporter we assume that the high affinity transport (up_2_) is reduced by 20%. For mutations in CEP or CK signalling we put the production of CEP or CK to zero. In Table 3 we show the length of the root system on the low nitrate side, the high nitrate side and their difference for wildtype as well as *nrt1.1*, *cep* and *ck* mutant plants after 6 days of simulated growth.

**Table 3.** Effect of *in silico* mutations on preferential root foraging.

| Plant | length low N side | length high N side | difference |
| --- | --- | --- | --- |
| WT | 36 | 128 | 92 |
| *nrt1.1* | 51 | 77 | 26 |
| *cep* | 39 | 96 | 57 |
| *ck* | 39 | 101 | 62 |

We see that all three mutants result in a significant reduction of differences in root length between low and high nitrate experiencing sides of the root, as one would expect for a decrease in preferential root foraging. Additionally, as can be expected from the suggested roles of CEP and CK signalling in the demand dependent enhancement of root growth on the high nitrate side, *cep* and *ck* mutants show less decrease in the reduction of root growth on the low nitrate side as compared to the *nrt1.1* mutant, which is involved in both repressing root growth for low nitrate and stimulating root growth for high nitrate. We do note that under our current model settings *cep* and *ck* mutants have a less strong effect on preferential foraging than the *nrt1.1* mutant. Importantly, the model we constructed is highly simplified and for example does not incorporate regulatory changes in transporter levels. Thus, an exact quantitative correspondence may not be reasonable to expect. Still, the fact that qualitatively the model reproduces all three mutants correctly strongly supports the validity of our model for unraveling root nitrate foraging.

## 4. Discussion

Preferential foraging of roots in nutrient rich soil patches is an important determinant of overall plant growth and fitness, enabling plants to survive in spatio-temporally varying conditions. Still, how exactly this preferential root foraging arises from the intricate network of internal and external nutrients sensing, signaling and subsequent responses has remained largely unclear. In the current study we investigated the case of preferential foraging for nitrate.

We used a highly simplified framework for modeling root system growth in which we incrementally incorporated the different known aspects of nitrate dependent root growth regulation. Specifically, we incorporated local nitrate level dependent growth stimulation and repression involving amongst other molecular mechanisms the nitrate dependent auxin transport of NRT1.1 (Krouk et al., 2010; Mounier et al., 2014), demand driven growth stimulation involving CEP signalling and NRT2.1 upregulation (Tabata et al., 2014; Ohkubo et al., 2017), as well as systemic nitrate status dependent survival, foraging and suppression responses (Giehl and Wirén, 2014). Additionally, we incorporated intra-root system competition for carbon allocation, as well as a CK mediated supply signalling that influences the efficiency of the CEP demand signalling. Our model correctly reproduces the strong reduction in preferential root foraging upon mutations in either NRT1.1, CEP signalling or CK signalling.

Our model outcomes suggest that preferential nitrate foraging does not merely involve demand and supply signaling. Instead, differences in root length occurring between split root plants experiencing heterogeneous nitrate levels (high on one low on the other side) and homogeneous high nitrate levels (so both sides high) also involve the (increased) engagement of systemic repression under homogeneous high nitrate conditions. Similarly, differences in root length occurring between plants experiencing heterogeneous nitrate levels and homogeneous low nitrate levels also involve the activation of foraging responses under homogeneous low nitrate conditions. Finally, our model indicates that competition for carbon resources may contribute to the asymmetry in root lengths occurring in split root experiments. Furthermore, our model suggests that the decrease in root growth at the low nitrate side in heterogeneous compared to homogeneous nitrate conditions does not require a supply signal from the high nitrate side, but rather arises from a reduced foraging response and an enhanced competition for carbon under heterogeneous conditions.

Based on the above, we suggest that in addition to comparing the high and low nitrate experiencing sides of a plant root system to roots experiencing on both sides high or on both sides low nitrate, comparisons should be extended to roots experiencing on both sides intermediate nitrate levels. We expect that this extension will help tease apart the effects of systemic repression, foraging from systemic demand and supply signaling effects. Additionally, by also applying this extension to the large scale transcriptomics analyses done recently to unravel key players in nitrate foraging (Ruffel et al., 2011; Canales et al., 2014; Poitout et al., 2018), we can gain more insight in which parts of the complex nitrate signaling and response network play a role in which of these different effects. Finally, given the potentially important role for carbon allocation in contributing to root system asymmetry, we suggest that experiments, in addition to reporting root system size, should also focus on potential differences and changes in shoot size.

A key outcome of our model is the suggested role for CK signaling in nitrate foraging. Previous studies have suggested a role for CK in systemic nitrate demand signaling (Ruffel et al., 2011; Poitout et al., 2018). This interpretation was based on the observed collapse of preferential root length increase on the high nitrate side in CK biosynthesis mutants, following the reasoning that differences in plants with one versus two high nitrate sides should arise from demand signaling. However, given the nitrate dependent production of CK, a role for CK in nitrate supply signalling appears more natural (Takei et al., 2002). Based and the observed effect of nitrate dependent, systemic CK signalling on the demand driven NRT2.1 upregulation (Poitout et al., 2018), we propose that CK signaling represents a supply signal that modulates the strength of CEP driven demand signaling. This hypothesis would imply that preferential foraging on the high nitrate side involves coordinated nitrate supply, demand and local presence signaling. Using our model we subsequently demonstrate that a supply signal impacting the efficiency of demand signaling would enable the plant to tune its extent of preferential foraging based on the balance between supply in demand. This helps prevent excessive investments in localized root growth and suppression of root growth elsewhere in situations where nitrate supply is highly restricted spatially and it is a more safe strategy for other parts of the root system to keep foraging for nitrate. To test our model prediction, we propose experiments in split root plants with a mildly high and a low nitrate side. If our predictions are correct, under normal conditions only moderate preferential foraging should occur, whereas after addition of CK this preferential foraging should increases because of the enhanced effectiveness of the demand signaling.

For future work, it will be interesting to implement our model within a spatially explicit, branching root architecture such as typically used in FSP models (for example (Schnepf et al., 2018)). Additionally, an interesting direction would be to attempt to extend the explanatory power of the model to the apparent sensitivity of root growth to temporal changes in nitrate levels (Shemesh et al., 2010a, 2010b, 2011). This likely requires the incorporation of additional model components, such as internal nitrate stores, or the regulation of expression of nitrate transporters (Aibara and Miwa, 2014)). Other interesting directions for expansion are to attempt to incorporate the effects of neighboring plants competing for nitrate (Ljubotina and Cahill Jr, 2019), requiring at least the incorporation of soil nitrate dynamics (Robinson, 2001), and potentially plant-plant communication mechanisms.

## Acknowledgements

We thank Bas van den Herik for helpful discussions.

## Author contributions

M.J.B. developed the initial version of the model and wrote an early version of the manuscript. J.S.T. analyzed model output, created figures and edited the manuscript. K.t.T. conceived the research and wrote the manuscript.

## References

Aibara, I., and Miwa, K. (2014). Strategies for Optimization of Mineral Nutrient Transport in Plants: Multilevel Regulation of Nutrient-Dependent Dynamics of Root Architecture and Transporter Activity. Plant Cell Physiol. 55, 2027–2036. doi:10.1093/pcp/pcu156.

Angle, J. S., Gross, C. M., Hill, R. L., and McIntosh, M. S. (1993). Soil Nitrate Concentrations under Corn as Affected by Tillage, Manure, and Fertilizer Applications. J. Environ. Qual. 147, 141–147. doi:10.2134/jeq1993.00472425002200010018x.

Araya, T., Miyamoto, M., Wibowo, J., Suzuki, A., Kojima, S., Tsuchiya, Y. N., et al. (2014). CLE-CLAVATA1 peptide-receptor signaling module regulates the expansion of plant root systems in a nitrogen-dependent manner. Proc Natl Acad Sci U S A. 111, 2029–2034. doi:10.1073/pnas.1319953111.

Canales, J., Moyano, T. C., Villarroel, E., and Gutiérrez, R. A. (2014). Systems analysis of transcriptome data provides new hypotheses about Arabidopsis root response to nitrate treatments. Front. Plant Sci. 5, 1–14. doi:10.3389/fpls.2014.00022.

Cerezo, M., Tillard, P., Filleur, S., Munos, S., Daniel-Vedele, F., and Gojon, A. (2001). Major Alterations of the Regulation of Root NO 3-Uptake Are Associated with the Mutation of Nrt2.1 and Nrt2.2 Genes in Arabidopsis. Plant Physiol. 127, 262–271. doi:10.1104/pp.127.1.262.

Crawford, N. M., and Glass, A. D. M. (1998). Molecular and physiological aspects of nitrate uptake in plants. Trends Plant Sci. 3, 389–395. doi:10.1016/S1360-1385(98)01311-9.

Fujikake, H., Yamazaki, A., Ohtake, N., Sueyoshi, K., Matsuhashi, S., Ito, T., et al. (2003). Quick and reversible inhibition of soybean root nodule growth by nitrate involves a decrease in sucrose supply to nodules. J. Exp. Bot. 54, 1379–1388. doi:10.1093/jxb/erg147.

Giehl, R. F. H., Gruber, B. D., and Wirén, N. Von (2014). It’s time to make changes: modulation of root system architecture by nutrient signals. J. Exp. Bot. 65, 769–778. doi:10.1093/jxb/ert421.

Giehl, R. F. H., and Wirén, N. Von (2014). Root Nutrient Foraging. Plant Physiol. 166, 509–517. doi:10.1104/pp.114.245225.

Girin, T., El-Kafafi, E.-S., Widiez, T., Erban, A., Hubberten, H.-M., Kopka, J., et al. (2010). Identification of Arabidopsis Mutants Impaired in the Systemic Regulation of Root Nitrate Uptake by the Nitrogen Status of the Plant. Plant Physiol. 153, 1250–1260. doi:10.1104/pp.110.157354.

Gruber, B. D., Giehl, R. F. H., Friedel, S., and Wirén, N. Von (2013). Plasticity of the Arabidopsis Root System under Nutrient Deficiencies. Plant Physiol. 163, 161–179. doi:10.1104/pp.113.218453.

Guan, P. (2017). Dancing with Hormones: A Current Perspective of Nitrate Signaling and Regulation in Arabidopsis. Front. Plant Sci. 8, 1–20. doi:10.3389/fpls.2017.01697.

Guan, P., Wang, R., Nacry, P., Breton, G., Kay, S. A., Pruneda-paz, J. L., et al. (2014). Nitrate foraging by Arabidopsis roots is mediated by the transcription factor TCP20 through the systemic signaling pathway. Proc. Natl. Acad. Sci. U.S.A. 111, 15267–15272. doi:10.1073/pnas.1411375111.

Hellebrand, H. J., Scholz, V., Kern, J., and Kavdir, Y. (2005). N2O Release During Cultivation of Energy Crops. Agrar. Forsch. 11, 114–124.

Jin, Z., Zhu, Y., Li, X., Dong, Y., and An, Z. (2015). Soil N retention and nitrate leaching in three types of dunes in the Mu Us desert of China. Sci. Rep. 5, 1–8. doi:10.1038/srep14222.

Krouk, G., Lacombe, B., Bielach, A., Perrine-Walker, F., Malinska, K., Mounier, E., et al. (2010). Nitrate-Regulated Auxin Transport by NRT1.1 Defines a Mechanism for Nutrien Sensing in Plants. Dev. Cell 18, 927–937. doi:10.1016/j.devcel.2010.05.008.

Lally, D., Ingmire, P., Tong, H.-Y., and He, Z.-H. (2001). Antisense Expression of a Cell Wall-Associated Protein Kinase, WAK4, Inhibits Cell Elongation and Alters Morphology. Plant Cell 13, 1317–1331. doi:10.1105/tpc.13.6.1317.

Little, D. Y., Rao, H., Oliva, S., Daniel-Vedele, F., Krapp, A., and Malamy, J. E. (2005). The putative high-affinity nitrate transporter NRT2.1 represses lateral root initiation in response to nutritional cues. Proc. Natl. Acad. Sci. 102, 13693–13698. doi:10.1073/pnas.0504219102.

Ljubotina, M. K., and Cahill Jr, J. F. (2019). Effects of neighbour location and nutrient distributions on root foraging behaviour of the common sunflower. Proc. R. Soc. B 286. doi:10.1098/rspb.2019.0955.

Ma, W., Li, J., Qu, B., He, X., Zhao, X., Li, B., et al. (2014). Auxin biosynthetic gene TAR2 is involved in low nitrogen-mediated reprogramming of root architecture in Arabidopsis. Plant J. 78, 70–79. doi:10.1111/tpj.12448.

Marcelis, L. F. M. (1996). Sink strength as a determinant of dry matter partitioning in the whole plant. J. Exp. Bot. 47, 1281–1291. doi:10.1093/jxb/47.Special_Issue.1281.

Mounier, E., Pervent, M., Ljung, K., Gojon, A., and Nacry, P. (2014). Auxin-mediated nitrate signalling by NRT1.1 participates in the adaptive response of Arabidopsis root architecture to the spatial heterogeneity of nitrate availability. Plant, Cell Environ. 37, 162–174. doi:10.1111/pce.12143.

Naz, M., Luo, B., Guo, X., Chen, J., and Fan, X. (2019). Overexpression of Nitrate Transporter OsNRT2.1 Enhances Nitrate-Dependent Root Elongation. Genes (Basel). 10, 1–18. doi:10.3390/genes10040290.

Ohkubo, Y., Tanaka, M., Tabata, R., Ogawa-Ohnishi, M., and Matsubayashi, Y. (2017). Shoot-to-root mobile polypeptides involved in systemic regulation of nitrogen acquisition. Nat. Plants 3, 1–6. doi:10.1038/nplants.2017.29.

Okamoto, Y., Suzuki, T., Sugiura, D., Kiba, T., Sakakibara, H., and Hachiya, T. (2019). Shoot nitrate underlies a perception of nitrogen satiety to trigger local and systemic signaling cascades in Arabidopsis thaliana. Soil Sci. Plant Nutr. 65, 56–64. doi:10.1080/00380768.2018.1537643.

Poitout, A., Crabos, A., Petřík, I., Novák, O., Krouk, G., Lacombe, B., et al. (2018). Responses to Systemic Nitrogen Signaling in Arabidopsis Roots Involve trans-Zeatin in Shoots. Plant Cell 30, 1243–1257. doi:10.1105/tpc.18.00011.

Remans, T., Nacry, P., Pervent, M., Filleur, S., Diatloff, E., Mounier, E., et al. (2006). The Arabidopsis NRT1.1 transporter participates in the signaling pathway triggering root colonization of nitrate-rich patches. Proc. Natl. Acad. Sci. 103, 19206–19211. doi:10.1073/pnas.0605275103.

Robinson, D. (2001). Root proliferation, nitrate inflow and their carbon costs during nitrogen capture by competing plants in patchy soil. Plant Soil 232, 41–50. doi:10.1023/A:101377818094.

Ruffel, S., Krouk, G., Ristova, D., Shasha, D., Birnbaum, K. D., and Coruzzi, G. M. (2011). Nitrogen economics of root foraging: Transitive closure of the nitrate-cytokinin relay and distinct systemic signaling for N supply vs. demand. Proc. Natl. Acad. Sci. U. S. A. 108, 18524–18529. doi:10.1073/pnas.1108684108.

Sasse, J., Jordan, J. S., DeRaad, M., Whiting, K., Zhalnina, K., and Northen, T. (2019). Root morphology and exudate availability is shaped by particle size and chemistry in Brachypodium distachyon. bioRxiv. doi:10.1101/651570.

Schnepf, A., Leitner, D., Landl, M., Lobet, G., Mai, T. H., Morandage, S., et al. (2018). CRootBox: a structural-functional modelling framework for root systems. Ann. Bot. 121, 1033–1053. doi:10.1093/aob/mcx221.

Shemesh, H., Arbiv, A., Gersani, M., Ovadia, O., and Novoplansky, A. (2010a). The Effects of Nutrient Dynamics on Root Patch Choice. PLoS One 5, 1–6. doi:10.1371/journal.pone.0010824.

Shemesh, H., Ovadia, O., and Novoplansky, A. (2010b). Anticipating future conditions via trajectory sensitivity. Plant Signal. Behav. 5, 1501–1503. doi:10.4161/PSB5.11.13660.

Shemesh, H., Rosen, R., Eshel, G., Novoplansky, A., and Ovadia, O. (2011). The effect of steepness of temporal resource gradients on spatial root allocation. Plant Signal. Behav. 6, 1356–1360. doi:10.6141/psb.6.9.16444.

Sun, H., Tao, J., Bi, Y., Hou, M., Lou, J., Chen, X., et al. (2018). OsPIN1b is Involved in Rice Seminal Root Elongation by Regulating Root Apical Meristem Activity in Response to Low Nitrogen and Phosphate. Sci. Rep. 8, 1–11. doi:10.1038/s41598-018-29784-x.

Tabata, R., Sumida, K., Yoshii, T., Ohyama, K., Shinohara, H., and Matsubayashi, Y. (2014). Perception of root-derived peptides by shoot LRR-RKs mediates systemic N-demand signaling. Science (80-.). 346, 343–346. doi:10.1126/science.1257800.

Takei, K., Takahashi, T., Sugiyama, T., Yamaya, T., and Sakakibara, H. (2002). Multiple routes communicating nitrogen availability from roots to shoots: a signal transduction pathway mediated by cytokinin. J. Exp. Bot. 53, 971–977. doi:10.1093/jexbot/53.370.971.

Taylor, A. A., De-Felice, J., and Havill, D. C. (1982). Seasonal Variation In Nitrogen Availability And Utilization In An Acidic And Calcareous Soil. New Phytol. 92, 141–152. doi:10.1111/j.1469-8137.1982.tb03370.x.

Terasaka, K., Blakeslee, J. J., Titapiwatanakun, B., Peer, W. A., Bandyopadhyay, A., Makam, S. N., et al. (2005). PGP4, an ATP Binding Cassette P-Glycoprotein, Catalyzes Auxin Transport in Arabidopsis thaliana Roots. Plant Cell 17, 2922–2939. doi:10.1105/tpc.105.035816.

van de Sande-Bakhuyzen, H. L. (1928). Studies Upon Wheat Grown Under Constant Conditions. Plant Physiol. 3, 7–30. doi:10.1104/pp.3.1.1.

Vidal, E. A., Álvarez, J. M., and Gutiérrez, R. A. (2014). Nitrate regulation of AFB3 and NAC4 gene expression in Arabidopsis roots depends on NRT1.1 nitrate transport function. Plant Signal. Behav. 9, 1–4. doi:10.4161/psb.28501.

Vidal, E. A., Araus, V., Lu, C., Parry, G., Green, P. J., Coruzzi, G. M., et al. (2010). Nitrate-responsive miR393/AFB3 regulatory module controls root system architecture in Arabidopsis thaliana. Proc. Natl. Acad. Sci. 107, 4477–4482. doi:10.1073/pnas.0909571107.

Vidal, E. A., Moyano, T. C., Riveras, E., Contreras-López, O., and Gutiérrez, R. A. (2013). Systems approaches map regulatory networks downstream of the auxin receptor AFB3 in the nitrate response of Arabidopsis thaliana roots. Proc. Natl. Acad. Sci. 110, 12840–12845. doi:10.1073/pnas.1310937110/.

Weil, R. R., and Brady, N. C. (1996). The Nature and Properties of Soil. Fiftheenth.

Xu, P., and Cai, W. (2019). Nitrate-responsive OBP4-XTH9 regulatory module controls lateral root development in Arabidopsis thaliana. PLoS Genet. 15, 1–27. doi:10.1371/journal.pgen.1008465.

